# Second-generation dual-channel visible light optical coherence tomography enables wide-field, full-range, and shot-noise limited retinal imaging

**DOI:** 10.1101/2022.10.05.511048

**Authors:** Jingyu Wang, Stephanie Nolen, Weiye Song, Wenjun Shao, Wei Yi, Ji Yi

**Affiliations:** Department of Ophthalmology, Johns Hopkins University, Baltimore USA; Department of Biomedical Engineering, Johns Hopkins University, Baltimore USA; Department of Medicine, Boston University School of Medicine, Boston Medical Center, Boston, USA; Department of Mechanical Engineering, Shandong University, Ji Nan, China

## Abstract

Visible light optical coherence tomography (VIS-OCT) is an emerging ophthalmic imaging method uniquely featured by ultrahigh depth resolution, retinal microvascular oximetry, and distinct scattering contrast in the visible spectral range. However, the clinical utility of VIS-OCT is impeded by the fundamental trade-off between the imaging depth range and axial resolution, determined by the spectral resolution and bandwidth respectively. While the full potential of VIS-OCT is leveraged by a broad bandwidth, the imaging depth is inversely sacrificed. The effective depth range is further limited by the wavelength-dependent roll-off that the signal-to-noise ratio (SNR) reduces in the deeper imaging range, more so in shorter wavelength. To address this trade-off, we developed a second-generation dual-channel VIS-OCT system including the first *linear-in-k* VIS-OCT spectrometer, reference pathlength modulation, and per A-line noise cancellation. All combined, we have achieved 7.2dB roll-off over the full 1.74 mm depth range (water) with shot-noise limited performance. The system uniquely enables >60° wide-field imaging over large retinal curvature at peripheral retina and optic nerve head, as well as high-definition imaging at ultrahigh 1.3 um depth resolution (water). The dual-channel design includes a conventional near infrared (NIR) channel, compatible with Doppler OCT and OCT angiography (OCTA). The comprehensive structure-function measurement by 2^nd^-Gen VIS-OCT system is a significant advance towards broader adaptation of VIS-OCT in clinical applications.

## 1 Introduction

Since its conception in the 1990s, optical coherence tomography (OCT) has been one of the most influential imaging methods in Biophotonics, and revolutionized the clinical practice in ophthalmology [1]. Visible light optical coherence tomography (VIS-OCT) is a burgeoning modification of OCT that uses visible instead of near-infrared (NIR) illumination and has been demonstrated for preclinical and clinical ophthalmic imaging [2–4]. One advantage of VIS-OCT is that visible wavelengths often demonstrate higher levels of backscattering from biological tissues than longer near-infrared wavelengths [5], and reveal distinct spectral contrast in imaging pigmentation in the eye (e.g. rhodopsin in photoreceptors, melanin [6,7], hemoglobin [8–10]). More importantly, the shorter central wavelengths of visible light spectra result in improved transverse resolution and, when bandwidth is maintained, improved axial resolution [11]. This improved resolution allows detailed imaging of structures such as Bruch’s membrane [12,13], sub-bands in the outer segment of photoreceptors [14], sub-layers of inner plexiform layer [15], and the texture of nerve fibers [13]. Another key application of VIS-OCT is microvascular oximetry in the retina, leveraging the strong oxygen-dependent absorption spectrum of hemoglobin in the visible range [16,17]. By performing 3D segmentation and spatio-spectral analysis, VIS-OCT specifically extracts signals within blood vessels to calculate the blood oxygen saturation [8,18,19] and further oxygen metabolic rate when incorporating Doppler blood flow measurements [19].

However, as with all budding technologies, there are still challenges in VIS-OCT that must be addressed for broader clinical utilities. The most fundamental challenge arises from the trade-off between imaging depth range and axial resolution. Current VIS-OCT systems use spectral domain or Fourier domain configurations with a linear line scan camera to record the interferogram. While the full potential of VIS-OCT prefers a broader bandwidth to achieve *e.g*., a wider spectral contrast and better axial resolution, the spectral resolution (unit: nm/pixel) is proportionally enlarged and thus reducing the total imaging depth range. The limited depth range is further exacerbated by the sensitivity roll-off over depth [20], a common characteristic in all FDOCT systems. The roll-off is particularly problematic in broadband FDOCT because of the non-linear distribution of wavenumbers over the pixels [21], resulting in wavelength-dependent roll-off that compromises the total performance [20,22,23]. To improve spectral resolution and imaging depth, line cameras with high pixel numbers can be used [24], with the caveat of slower imaging speed and challenging optical design to account for aberration for a long linear pixel array. The limited depth range is particularly cumbersome at the presence of involuntary eye movement during imaging, as well as wide-field imaging outside macula with large retina curvatures.

To overcome this fundamental trade-off in VIS-OCT, we have developed a second-generation device that incorporated several enabling designs. First, to overcome the wavelength-dependent roll-off, we developed the first linear-in-*k* VIS-OCT spectrometer covering 140nm bandwidth (500-640nm). Second, we implemented a reference pathlength modulation to achieve full-range OCT, doubling the total imaging depth range, as well as to correct retinal curvature for wide-field imaging. Third, to eliminate the excessive noise at the zero-delay line due to the supercontinuum generation, we implemented per-A-line noise cancellation in the visible channel to achieve shot-noise limited imaging.

By the above designs, we have achieved in total 7.2dB roll-off (50% of the state-of-the-art VIS-OCT spectrometer) over a full range of 1.74 mm imaging depth (water), with a depth resolution down to 1.3 μm (water). We demonstrated a wide-field VIS-OCT retinal imaging >60° viewing angle in retina, largest so far to the best of our knowledge; a full-range circular scanning at optic nerve head (ONH); OCT angiography (OCTA) by NIR-OCT channel; and finally high-definition ultrahigh resolution retinal imaging. Human imaging interaction with the device is improved by minimizing visible light exposure through a rapid line rate of 120 kHz. The system design is a significant step forward to improve the clinical utility, and the imaging capability could potentially generate impactful insight by revealing structural and functional changes in retinal and/or neurodegenerative pathologies.

## 2 Result

### 2.1 Overview of system design

The configuration of the 2^nd^ Gen dual-channel VIS-OCT is shown in Fig. **1a**. Dual-channel imaging was implemented by two wavelength-division-multiplexers (WDMs) to combine and split the visible and NIR bands, and an ultra-broadband fiber coupler forming a Michelson interferometer. Two channels can operate alone or simultaneously, and complement each other where VIS-OCT can provide ultrahigh resolution imaging, retinal oximetry, while NIR-OCT is used for initial alignment, OCT angiography (OCTA), and Doppler OCT. The dual-channel broadband light is provided by a supercontinuum source with two filter sets to output visible band (500-650nm) and NIR band (750-900nm) respectively. To account for the chromatic shift over the large bandwidth, a custom-designed achromatizing lens (AL) [25] was installed in the sample arm. In addition, a tunable lens was used for focal adjustment. The detailed components and optics are provided in the method section. The depth resolution is 1.3 and 3.7 microns in water, and the power at cornea is <0.22 and 0.9 mW in VIS-OCT and NIR-OCT, respectively. All data shown here were acquired at 120 kHz Aline rate.

**Fig. 1.**
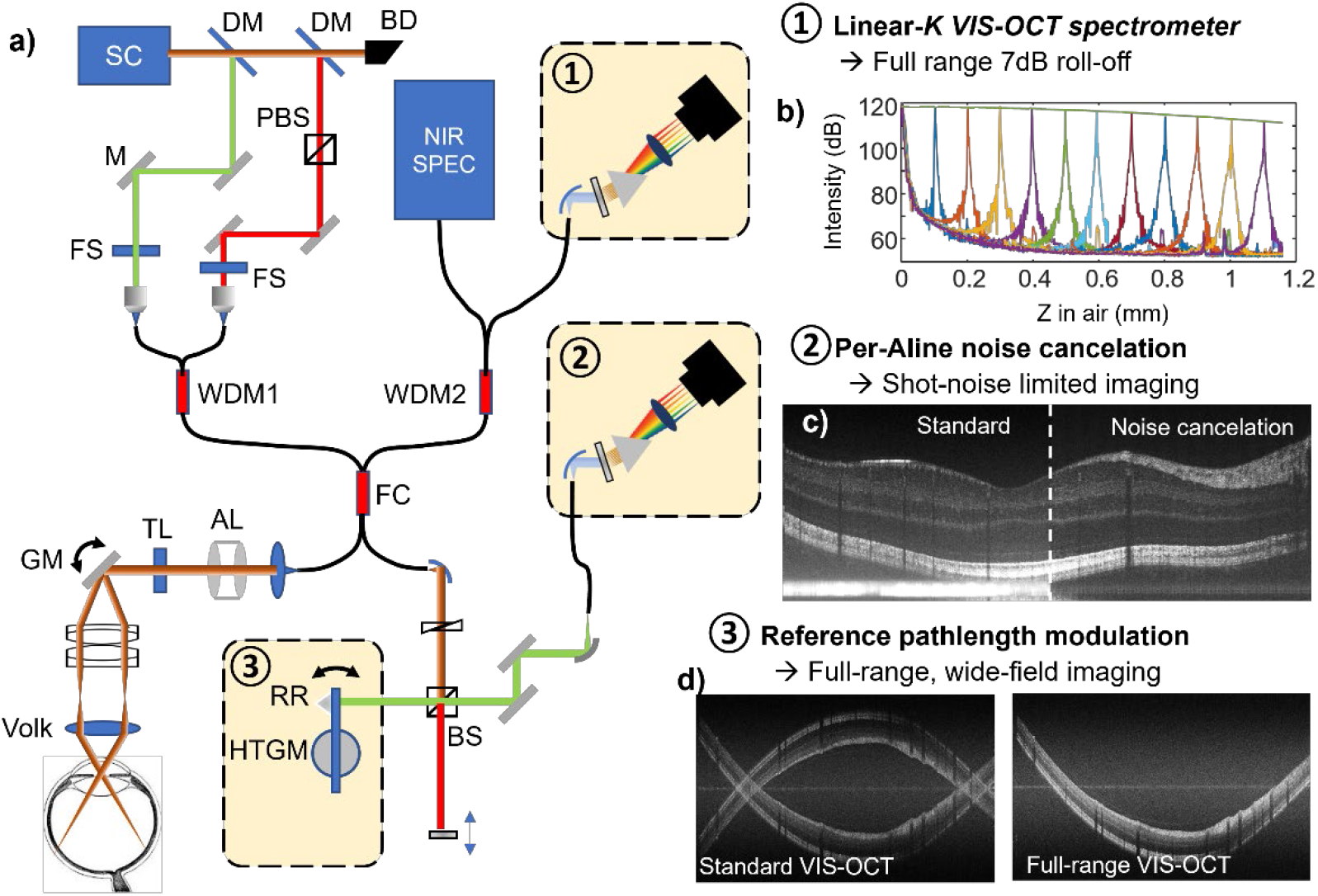
Overview of 2^nd^ generation dual-channel VIS-OCT. Left: (a) schematic of the optomechanical configuration. Three key enabling designs for full-range, wide-field imaging are 1) linear-in-k VIS-OCT spectrometer; 2) per-Aline noise cancelation by a second spectrometer; and 3) reference pathlength modulation. Right: summary of the imaging performance by three optical designs, achieving b) 7.2dB roll-off over the entire imaging range; c) short-noise limited performance; and d) doubling the imaging depth range.

The major effort of this paper is to overcome the trade-off between the spectral bandwidth, axial resolution, and imaging depth in VIS-OCT. To this end, three key design components are highlighted in Fig. 1 snapshots: 1) *Linear-in-K* spectrometers that eliminate the wavelength-dependent roll-off (**Fig. 1b**); 2) Reference pathlength modulation that achieves full range imaging, double the imaging depth while maintaining the same resolution (**Fig. 1c**); and 3) Noise cancellation to suppress the excessive noise to enable high-contrast and shot-noise limited imaging (**Fig. 1d**). The corresponding components are highlighted in Fig. 1a, and explained in detail in following sections.

**Fig. 2.**
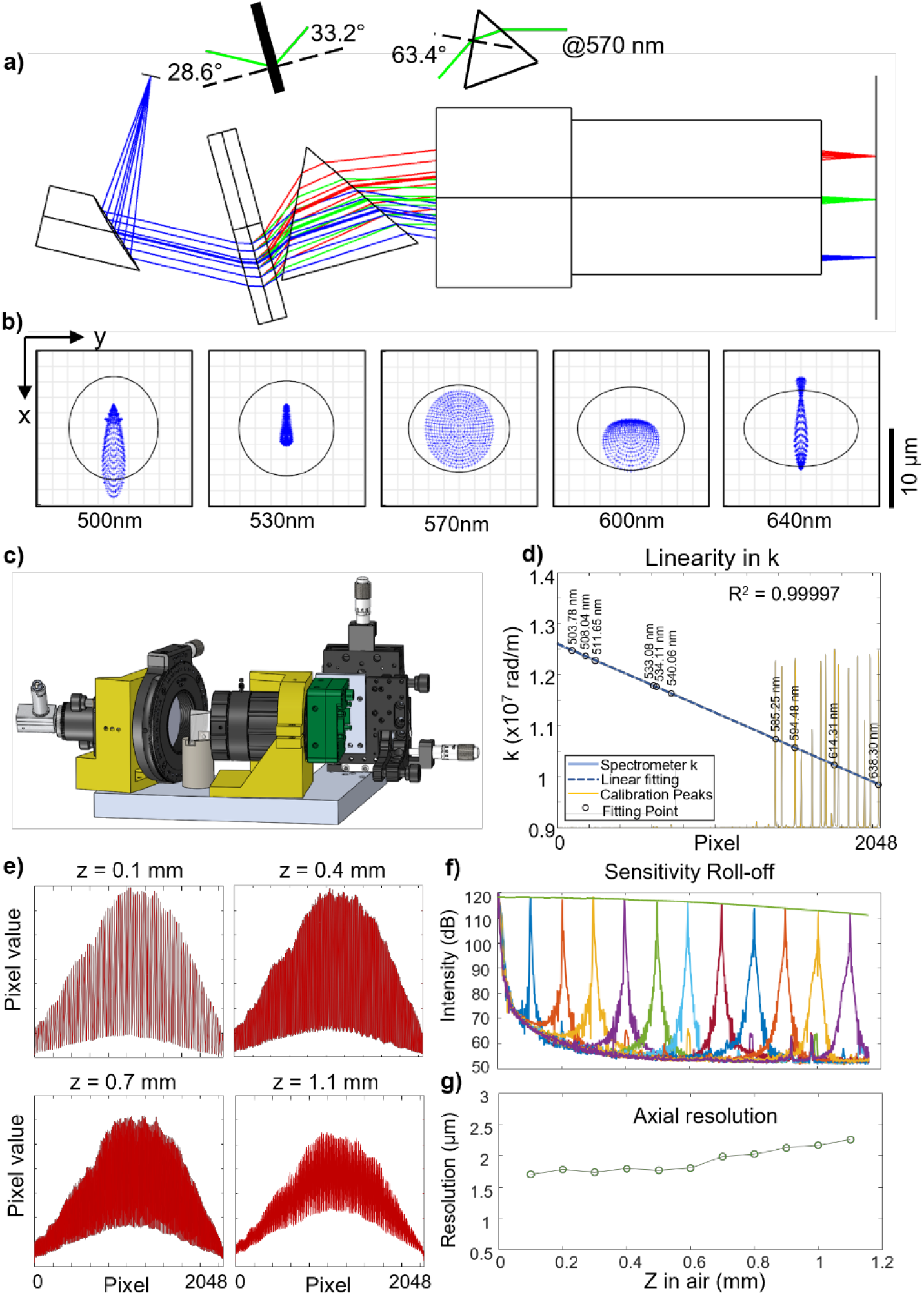
Design and characterization of linear-in-k VIS-OCT spectrometer. (a-c) The optical layout, the scatter plots in Zemax at different wavelength over the spectral range, and the 3D model of the spectrometer. Three colors in (a) traced 500, 570, and 640 nm respectively. (d) Spectrometer calibration using Mercury-Neon lamp, and the corresponding characteristic peaks. Pixel index is highly linear to the wave number, *k* by R^2^=0.99997. (e) The raw interferogram spectra by two mirror reflections with varying optical delay. (f-g) Sensitive roll-off and axial resolution characterization over the entire imaging depth range.

### 2.2 Linear-in-*k* VIS-OCT spectrometer

The design principle of linear-in-*k* spectrometer has been described in the literature [22], and first achieved here for VIS-OCT. Figure **2a** shows the Zemax model, consisting of a collimator (*f*=50mm), a transmission grating (Wasatch Photonics, 1800 lines/mm), an equilateral dispersive prism (Thorlabs, F2), a focusing lens (Edmundoptics, CA series, *f*=75mm), and a line scan camera (e2V, Octoplus 2048, 250kHz). The prism introduces a nonlinear dispersion that resulted in a linear-in-*k* relation when the light was focused on the camera pixels. We performed a Zemax simulation on the scatter plots at 5 wavelengths, showing that the diffraction-limited resolutions were achieved (Fig. **2b**). The 3D rendering model included all the mechanical components in Fig. **2c**. The mountings in yellow were 3D-printed, while the line scan camera was mounted on a 5-axis stage with x-y-z translation and yaw-pitch angle adjustment for precise alignment of the camera sensor. We calibrated the spectrometer using a Mercury lamp and confirmed excellent linearity (R^2^>0.999) in wave number k = 2π/λ, where λ is the wavelength (**Fig. 2d**). The exact wavelength coverage is from 498.9 to 639.3 nm. We next evaluated the roll-off performance of the linear-in-*k* spectrometer using a reflective mirror to provide the sample signal. The interferogram fringe amplitudes at varying depths are shown in Fig. **2e**. The contrast of the fringe decreased with the increasing depth, yet the change is consistent over the entire bandwidth (*i.e*. independent of wavelength), a distinct benefit of the linear-in-*k* spectrometer. The fringe contrast is slightly tapered off at the edge of the bandwidth at the deeper range, presumably due to the field curvature of the focusing lens. We characterized the overall roll-off performance in Fig. **2f** and measured 7.2 dB decay by the polynomial fitting over the entire depth range of 1.16 mm in air, and 0.9 mm in water. The roll-off is ~50% that of the state-of-the-art conventional VIS-OCT spectrometer, where the total roll-off over the imaging range is estimated at ~14-15dB [20,24]. Full-width-half-maximum (FWHM) width from the mirror surface was used to characterize the axial resolution in Fig. **2g**, down to ~1.7 μm (air) and 1.3 μm (water) close to the DC delay line. The resolution is maintained within 2.3 μm (air) and 1.7 μm (water) through the entire depth range.

### 2.3 Per A-line noise cancellation by simultaneous reference signal collection

All current VIS-OCT systems used supercontinuum (SC) light sources. However, the nonlinear phenomena that occur when these sources generate light lead to power fluctuations in the spectra [26], known as relative intensity noise (RIN). For SC sources, the RIN is the main contributor to excess noise [27], which is the noise that surpasses the shot (Poisson) noise caused by the quantum nature of photons [28]. Noise can be reduced by increasing the spectrometer exposure time, essentially averaging out the RIN fluctuations [28] but will also increase imaging time. Another noise minimizing technique is to use a balanced detection method [27,29–31]. By leveraging the temporal signature of RIN and calibrating two spectrometers with sub-pixel precision [32], recent work showed that the RIN background can be effectively removed [33,31].

We implemented noise cancellation by simply splitting the reference signal to a second identical VIS-OCT spectrometer (Fig. 1a, highlighted box 2). Two spectrometers were triggered simultaneously, acquiring temporal spectral signals, *x*_1_(*n, T*), *x*_2_(*p, T*), where n, p = 1-2048 denotes camera pixel index, and *t* denotes time. Continuous acquisition allows calculating pixel-wise temporal correlation between pixels in two spectrometers, shown in Fig. **3a**. The maximum values of correlation coefficients were used for pixel calibrations between two spectrometers, resulting in a sub-pixel alignment precision (Refer to section 4.3 for detail processes) and a close spectral matching after calibration/scaling (Fig. **3b**).

**Fig. 3.**
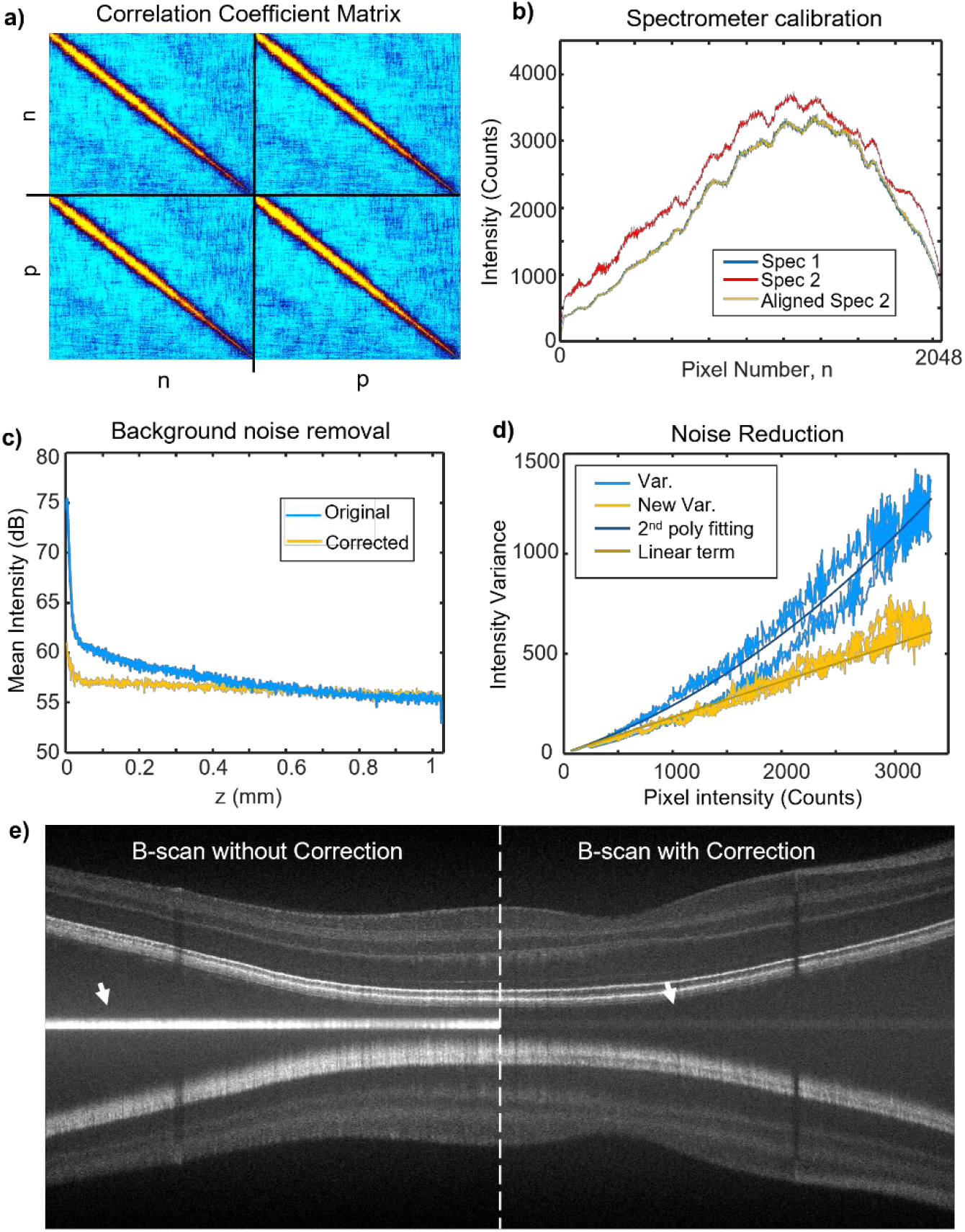
Per A-line excessive noise cancellation. (a) Pixel-pixel correlation matrix over continuous acquisition between two VIS-OCT spectrometers. Top-left and bottom-right are auto-correlation matrix for each spectrometer. Top-right and bottom-left are cross-correlation between two spectrometers (b) Spectrum match before and after spectrometer alignment. (c) Background noise in spatial domain before and after noise cancellation. (d) Intensity variance per intensity value over all pixels before and after noise cancellation. (e) Example of B-scans with and w/o noise cancellation.

We took 512 acquisitions of *x*_1_(*n, T* = 1 − 512) and generated a conventional background B-scan image. The DC components were removed by subtracting the mean spectrum 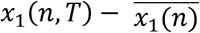. The averaged A-line signal is then shown in Fig. **3c**, where RIN produced a large noise background at zero-delay depth. In comparison, by per A-line noise cancelation, we removed the DC components by *x*_1_(*n, T*) – *x*_2_′(*n, T*), where *x*_2_′(*n, T*) is the calibrated/scaled spectra from the second spectrometer. The averaged A-line over the same 512 acquisitions reduces the background at the zero-delay depth by 4 folds, as well as in the shallower imaging depth. We further characterized the noise statistics by plotting the temporal variance σ^2^ for all 2048 pixels versus their mean pixel value (blue curve in Fig. **3d**). For the original *x*_1_(*n, T*), a significant quadratic term is present when fitted the data with a second order polynomial, representing the RIN [34]. We then plotted the variance for *x*_1_(*n,T*) – *x*_2_′(*n,T*), versus the same mean pixel values (yellow curve in Fig. **3d**). It matches well with the linear term from the polynomial fitting, representing a Poisson distribution and a shot noise limited performance. The benefit of noise cancellation is clearly shown in an example VIS-OCT image in Fig. **3e**, where the bright background in the zero-delay depth is suppressed to a faint gray shadow as indicated by the white arrows. The noise cancellation is a critical step in achieving full-range VIS-OCT, since the zero-delay line lies in the middle of the image.

### 2.4 Wide-field and full-range VIS-OCT imaging

When studying regions outside of the macula, it is necessary to have sufficient imaging depth to account for the eye curvature. To address the trade-off between imaging resolution and depth, and to enable wide-field VIS-OCT imaging, we implemented reference pathlength modulations. The hardware setup was accomplished by mounting a retroreflector on a high torque galvanometer (HTGM) [35] depicted in Fig. **4a**. The rotation of the shaft would modulate a reference pathlength. For the fast scan modulation (Fig. 4b), the retroreflector moves slowly at a constant speed, introducing a phase modulation into the *k*-x domain within B-scan (Fig. 4c) where *x* denotes the fast axis. A Hilbert transform can be performed along x to obtain complex interferogram values [36]. This phase modulation removes the phase ambiguity when performing Fourier transform, so that the symmetric OCT image to the zero-delay line can be eliminated and the imaging range is doubled. Successful removal of the mirror image is evident when comparing a typical VIS-OCT image (Fig. **4d**) with full-range imaging (Fig. **4e**). Because we fully utilized the signal, the contrast and brightness in full-range VIS-OCT image is also improved compared with conventional VIS-OCT.

**Fig. 4.**
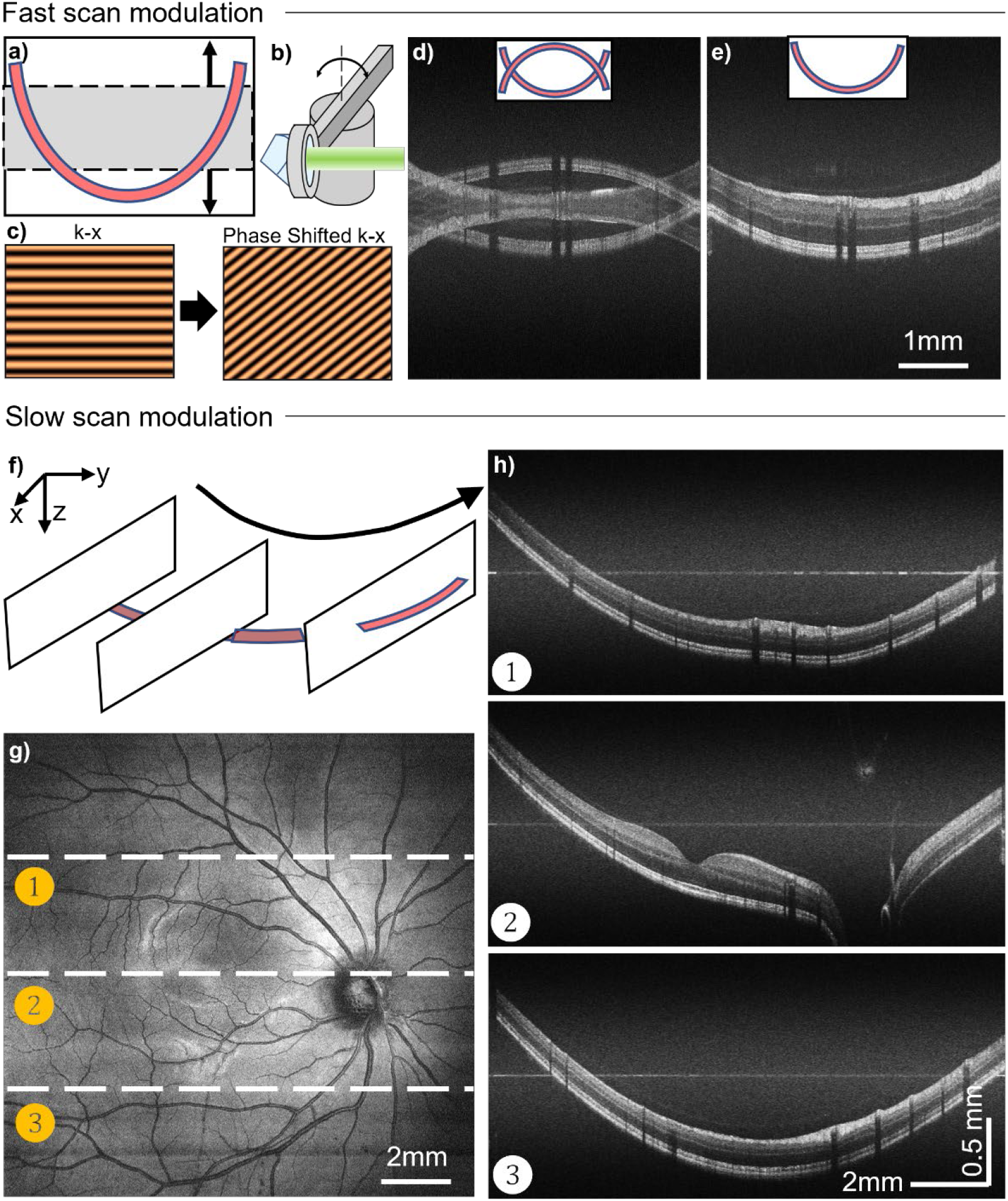
Full range and wide-field VIS-OCT enabled by the reference pathlength modulation. (a) Schematic model of the retroreflector (RR) and high-torque galvanometer (HTGM) to modulate the reference pathlength. (b-c) The phase introduced by path-length modulation to extend the imaging depth by fast scan modulation. (d-e) The comparison between conventional VIS-OCT and full range VIS-OCT to remove the Fourier transform ambiguity. (f) Schematic for slow scan modulation to compensate the retinal curvature by adjust the reference path per B-scan. (g-h) Wide-field VIS-OCT *en face* projection and the full-range VIS-OCT from three B-scans locations.

Within the same raster scanning, the slow scan modulation also permitted wide-field retinal imaging by moving the zero delay of the image in an arc corresponding to the eye curvature (Fig. 4f). The *en face* image in Fig. **4g** has corresponding cross sections in Fig. **4h** from three B-scan locations. The position of the retina is maintained in each full-range B-scan images, mitigating the typical displacement of the retina caused by the eye curvature. Meanwhile, the curvatures within each B-scan frames are entirely covered due to the extended imaging depth range (~1.7 mm in retina) (Fig. **4h**). The optics allows >60° viewing angle to image peripheral retina unattainable before (Supplemental Fig. 1).

The extended depth range is particularly advantageous in imaging optic nerve head (ONH) or peripheral retina, where large curvatures are present. We designed a disk scanning pattern where a series of dense circular B-scans (4096 Aline per B scan) were repeated with an increasing radius, covering a peripapillary disc area (Fig. **5a**). The expanded *en face* projection from VIS-OCT and NIR-OCT scans were shown in Fig. **5b** and **5c**, intersecting all major vessels in the inner retinal circulation branching and entering ONH. The dense circular scan can serve two purposes at the same time: *1*) to enable Doppler OCT, by calculating the phase delay between two adjacent A-lines in NIR OCT channel [37]; and *2*) to allow the fast scan modulation in VIS-OCT channel for full-range imaging. Figure **5d** and **5e** exemplified such circular scans in NIR-OCT and the corresponding phase contrast for Doppler OCT. We note that NIR channel is preferable for Doppler OCT due to the better penetration through vessel lumen than VIS-OCT for large vessels. Meanwhile, Fig. **5f** demonstrated the circular full-range VIS-OCT that the curvature of the parapapillary retina is well covered within the frame, which would otherwise be clipped.

**Fig. 5.**
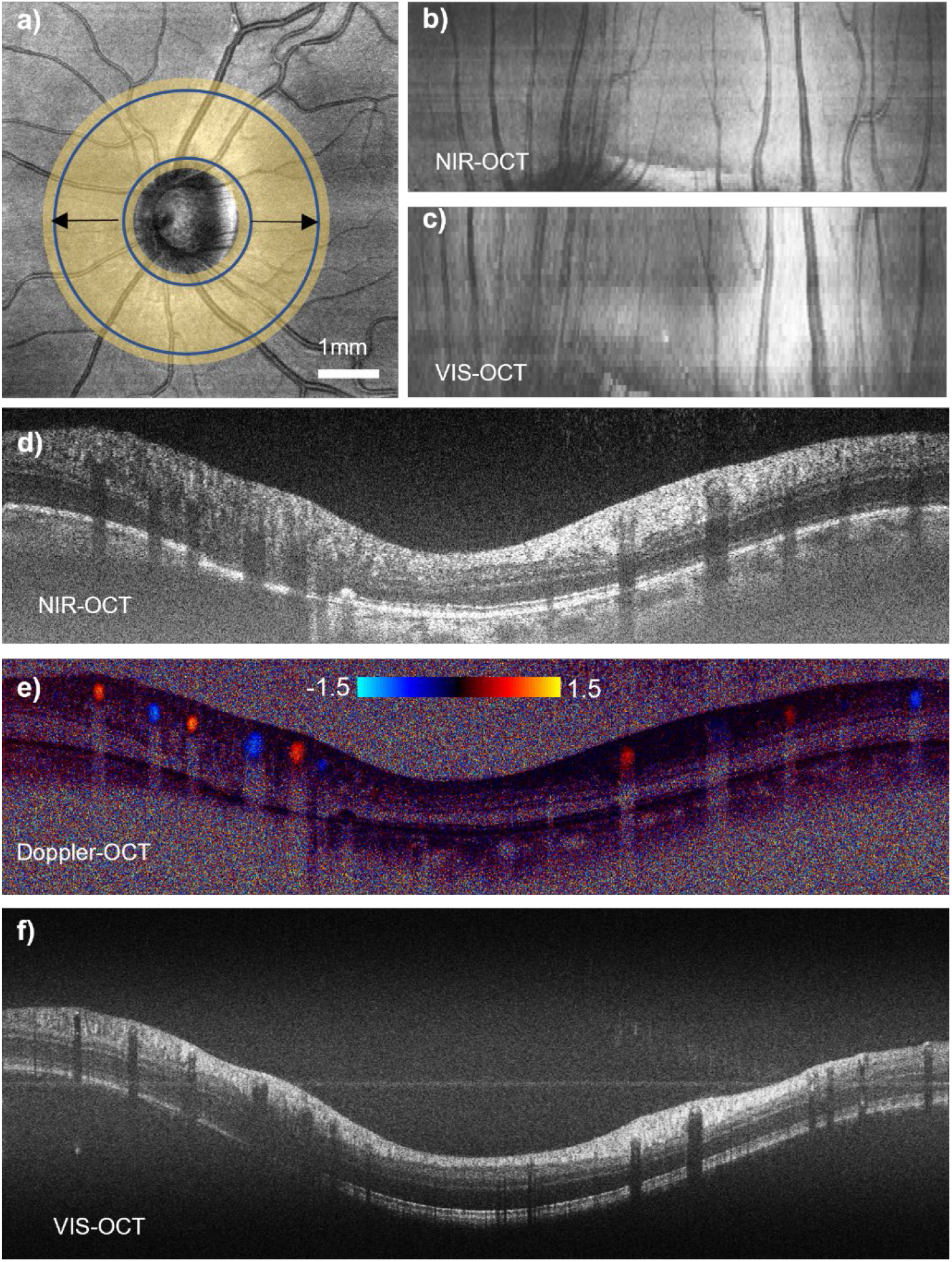
Full-range VIS-OCT imaging at optic nerve head (ONH). (a) *En face* projection of a VIS-OCT scan centered at ONH. Shaded area shows the disk area by circular scanning with increasing radius. (b-c) Expanded disk area samples all major vessels at peripapillary region. (d-f) Circular B-scan by NIR-OCT, Doppler OCT and VIS-OCT acquired simultaneously.

### 2.5 High-definition dual-channel structural imaging

By addressing the imaging range in VIS-OCT, we can fully leverage the high axial resolution enabled by the broadband linear-in-*k* VIS-OCT spectrometer. We adopted the speckle reduction protocol [38] and averaged 16 A-lines at each pixel to generate high-definition (HD) imaging. Figure **6a** and **6b** display the simultaneously acquired HD VIS-OCT and HD NIR-OCT images. The depth resolution in NIR channel is estimated to be ~3.7 microns in water based on the reference spectrum. The penetration into the choroid in VIS-OCT is weaker than NIR-OCT due to the stronger absorption by the retinal pigment epithelial (RPE) and choroidal pigment. The improvement of the resolution in HD VIS-OCT is visually apparent with finer anatomical stratification than HD NIR-OCT. We zoomed in on two areas in the outer segment of the photoreceptors and the inner plexiform layer (IPL) for a side-by-side comparison (Fig. **6c**-**6f**). HD VIS-OCT clearly resolved several bands beyond the outer segment of photoreceptor, including the IS/OS junction, cone outer segment tip (COST), rod outer segment tip (ROST), RPE and Bruch’s membrane (BM) [14] (Fig. **6c**). There are also five sub-bands resolvable in VIS-OCT in IPL, corresponding to different synaptic junctions between bipolar cells and retinal ganglion cells dendrites [3,15] (Fig. **6d**). When we zoomed-in on the nerve fiber layer (NFL) and ganglion cell layers (GCL), the fiber bundle texture is present in HD VIS-OCT, and a clear presentation of a thin GCL is evident (Fig. **6g**). Although the penetration into the deeper choroid is limited in VIS-OCT, we are still able to image the signal beyond choriocapillaris in the perifoveal and parafoveal areas (arrows in Fig. **6g**). Figure **6h** shows interesting structural stratification in the outer plexiform layer (OPL) where the photoreceptors synapse with bipolar cells. The deep capillary plexus in the retinal circulation is visible, noted by the hyperreflective signals above the OPL. We intentionally adjusted the contrast in Fig. **6h** to show the Henle’s fiber layer as a lighter band above the slightly brighter outer nuclear layer (ONL). Beyond the better anatomical imaging merely by the improved resolution in VIS-OCT, the signal contrast may also be distinct between VIS-OCT and NIR OCT due to the wavelength differences, suggesting potential scattering spectroscopic analysis [24,25,39].

**Fig. 6.**
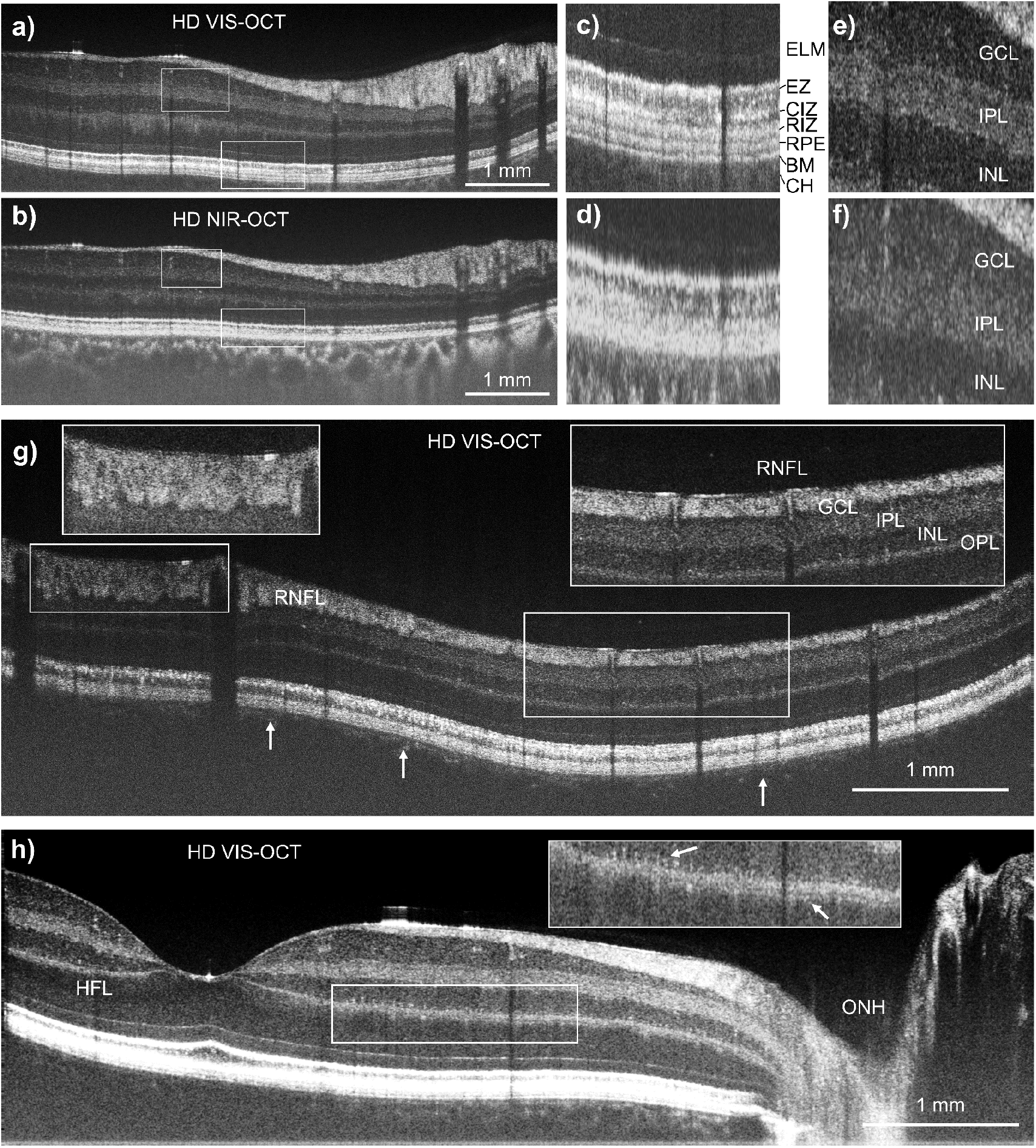
High-definition (HD) VIS-OCT reveals anatomical details with high resolution. (a-f) Comparison of VIS-OCT HD and NIR-OCT HD B-scan images, and Zoomed-in images showing anatomical layers resolvable by VIS-OCT HD. ELM: external limiting membrane; EZ: ellipsoid zone; CIZ: cone interdigitation zone; RIZ: rod interdigitation zone; RPE: retinal pigmented epithelium; BM: Bruch’s membrane; CH: choroid; GCL: ganglion cell layer; IPL: inner plexiform layer; INL: inner nuclear layer (g-h) VIS-OCT HD imaged RNFL texture, capillaries on OPL, as well as the sub-layers in OPL where photoreceptor synapses with bipolar cells. The contrast was adjusted in panel (h) to show the Henle’s layer in outer retina.

### 2.6 OCTA imaging

The dual-channel configuration in the device is compatible with clinical NIR-OCT and OCT angiography (OCTA). Figure 7 demonstrated an OCTA tiling overlaying on an *en face* image acquired by a wide filed NIR-OCT scan, covering a range of 13.3 mm × 7.4 mm. The angiogram was composed of five OCTA acquisitions, each with ~50% overlapping area with the adjacent patch, showing the major vessels as well as the single-capillary level network from internal limiting membrane (ILM) to outer plexiform layer (OPL).

**Fig. 7.**
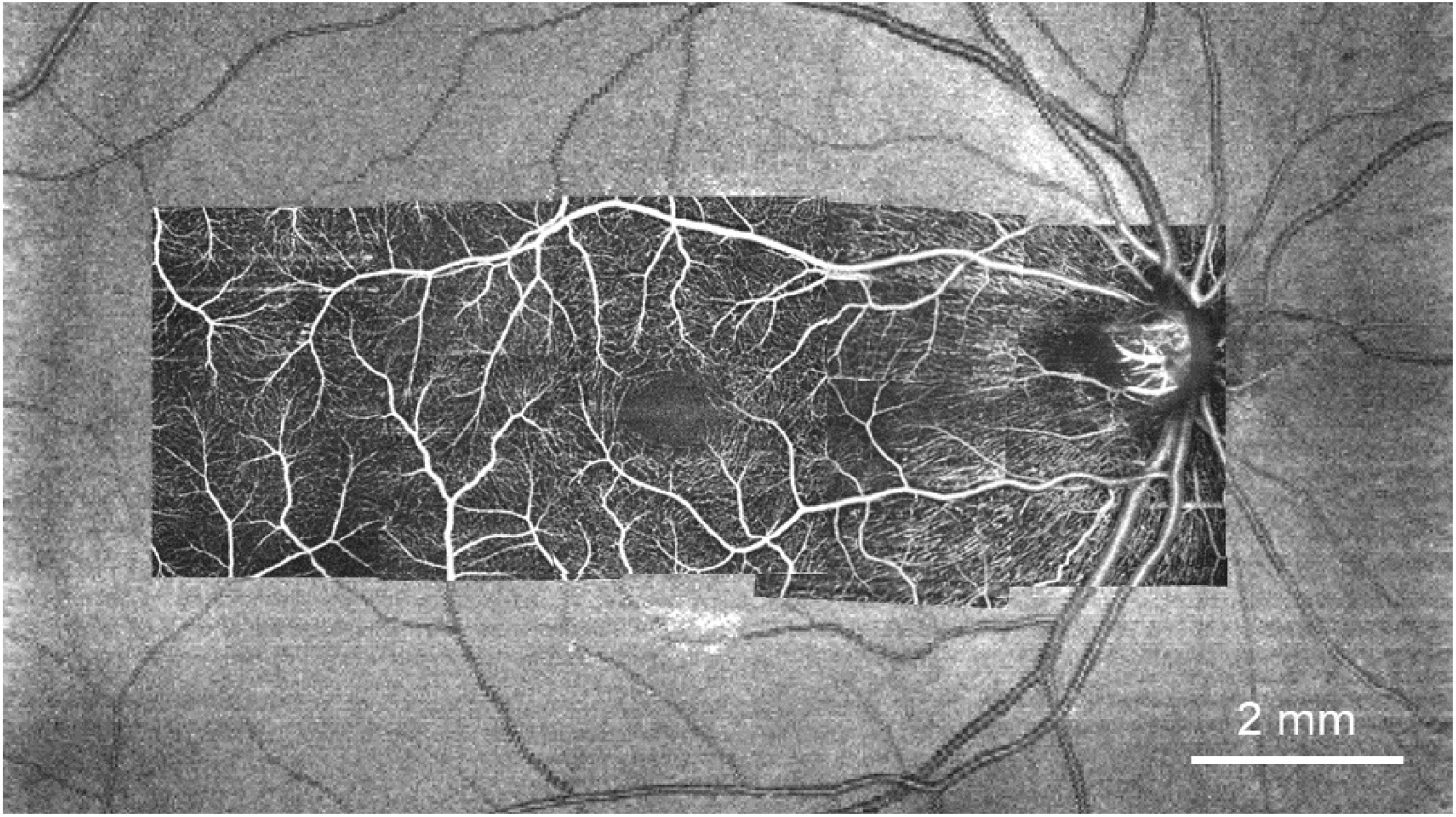
OCT angiography (OCT) by NIR channel showing microvasculature down to capillary level. The mosaic was generated from 5 acquisitions. Each OCTA image covers 3.3 by 3.3 mm FOV.

## 3 Discussion

In this paper, we presented the 2^nd^ Gen dual-channel VIS-OCT device that offers the best roll-off performance by far with a linear-in-*k* spectrometer, extended imaging range up to 1.8 mm in tissue at 1.3-1.7 microns depth resolution, and a wide-field imaging by the reference pathlength modulation. Implementing noise cancellation achieved shot-noise limited performance, a key step to remove the excessive noise near the zero-delay line for full-range VIS-OCT imaging. We showed that the dual-channel system not only offers the micron-level ultrahigh resolution VIS-OCT images and is also compatible with Doppler OCT and OCTA using the NIR channel. The practical advantage of NIR channel aided in rapid image alignment, a major practical challenge in operating VIS-OCT in clinics.

### 3.1 Further technical refinements

The spectrometer is the key component in VIS-OCT, given the stringent power safety limit and relatively smaller depth imaging range than NIR-OCT. The linear-in-*K* design eliminated the wavelength-dependent roll-off and significantly improved the overall imaging performance compared to the conventional design. The optical design enables diffraction-limited resolution, significantly smaller than the 10 μm pixel size in the camera, and thus the roll-off is determined solely by the pixel averaging. The additional refractive prism would introduce a slight light loss at two prism-air surfaces; however, the Fresnel refraction is polarization-dependent, which can be removed by adding a polarization controller before spectrometer. We also should note that the axial resolution in VIS-OCT can be further improved by tuning the spectral shape to be a top-hat instead of a semi-Gaussian, such that the spectrometer bandwidth can be fully utilized.

The noise cancellation in this presented form is a straightforward add-on in the existing Michelson interferometry, simply splitting and recording a fraction of the reference light. The previous methods used balance detection with a Mach–Zehnder interferometer as in a swept-source OCT system, subtracting two simultaneous interferogram with a phase shift of π to cancel out excessive noise and improve the signal to noise ratio [33,31]. While the balance-detection scheme still uses a line scan camera without gains, the doubled dynamic range offered by the additional camera does provide a signal boost. To this end, the balance-detection approach for per Aline noise cancellation is considered a better configuration. The full benefit of balance-detection in VIS-OCT would still need to wait for a swept source operating in visible light range.

### 3.2 Potential clinical applications

The reported device enables comprehensive capability to study structure/function relations in retina for a broad range of potential clinical applications. The ultrahigh resolution reveals structural details that is unattainable with conventional OCT imaging. The better layer characterization of nerve fiber layer, RGC, and sub-bands of IPL have clinical implications of optical neuropathy, such as glaucoma. In outer retina, while there are still debates on what anatomical layers in the outer segment of photoreceptors do those OCT bands correspond to [40–43], VIS-OCT images provide excellent depth resolutions to elucidate those debates and better imaging capabilities in visualize pathological changes in outer retina, such as macular degeneration, geographic atrophy, central serous retinopathy.

Another attractive potential of the system is the comprehensive analysis of perfusion function in the retina circulation systems, being able to combine measurement of blood oxygen saturation (sO_2_) [8], Doppler blood flow in major arterioles and venules [19], as well as OCTA at capillary level [2,9]. Retina, being the most metabolically active organ, heavily relies on microvascular system to maintain the functions. Perfusion dysfunction or abnormality is an important part of the pathogenesis in a broad range of clinical conditions, such as diabetic retinopathy, central retinal vein occlusion, choroid neovascularization, glaucoma *etc*. The measurement of oxygen consumptions either globally or locally may infer the metabolic function and can be correlated with structural changes.

By contrasting VIS-OCT and NIR-OCT images or simply within the broad visible spectrum, interesting spectroscopic information in retina can be potentially associated with different pathologies. A “true color” 3D reconstruction of mouse retina using RGB reconstruction in VIS-OCT was reported to examine the spectral contrast in photoreceptors, RPE, and the retinal lesions [39,44,45], where photo-sensing pigments and melanin plays important roles maintaining the physiological conditions. Similar analysis by VIS-OCT has performed in human retina, as well as by crossing the VIS-OCT and NIR-OCT images [4,24,25]. The spectroscopic analysis may reveal subtle structural contrasts determined by local refractive indices fluctuation, serving sensitive markers for detecting nanoscale changes beyond the resolution limit. For example, reflectance spectral analysis suggested cytoskeleton disruption in RGC axons, and has shown promising clinical applications in early glaucoma detection. The same paradigm can be applied in photoreceptors, potentially characterizing early cell damages.

### Conclusion

We have developed a 2^nd^ generation dual-channel VIS-OCT system to address the fundamental trade-off between image depth range and resolution and achieve the best roll-off performance so far for VIS-OCT (7.2dB over 1.8mm in retina), wide field-of-view (>60°), and shot-noise limited imaging. The comprehensive imaging capabilities by dual-channel design can be an important tool in studying structure/function relations in retinal pathologies.

## Acknowledgements

This study was supported by BrightFocus foundation (G2017077), and in part by NIH R01NS108464, R21029412, and R01EY032163.

## 4 Methods

### 4.1 System setup and control

The VIS (500 to 650 nm) and NIR (750 to 900 nm) illumination were from a supercontinuum light source (Superk EXTREME, EXU-OCT-6, NKT Photonics), initially selected by two dichroic mirrors (DMLP650R and DMLP900R, Thorlabs). Two bandpass filter sets in VIS and NIR channels further filtered the light (FEL0500+FES0650 VIS channel, and FEL0750+FES900 in NIR channel, Thorlabs). An electrical shutter (#87-208, Edmund Optics) was put in the VIS channel for alignment purposes. The filtered light of the dual-channel was coupled into the wavelength-division multiplexer (WDM 560/830, Customized, Thorlabs) to become the input of a 90/10 fiber coupler (FC, TW670R2A2, Thorlabs). The output end (10%) of the FC was connected to the sample arm and the other end (90%) was directed to the reference arm. In the sample arm, the light beam passed through a collimator (*f*=6mm, #63-714, Edmund Optics), a customized achromatizing lens [25], tunable lens (TL, EL-10-30-TC-VIS-12D, Optotune), two-axis galvanometer scanner (GVSM002, Thorlabs), and 3:1 telescope (*f*=75 mm, and *f*=25 mm, Volk 40D). The maximum light power on the pupil was controlled to be <0.22 mW (VIS) and 0.9 mW (NIR). In the reference arm, before the light enters a beam splitter (BS), it traverses a collimator (*f*=15mm, RC04APC-P01, Thorlabs) and a series of BK-7 dispersion plates in the optical path. The beam splitter worked with two edge filters (FES0700 and FEL0750, Thorlabs) to separate the VIS and NIR beam to a retroreflector (Edmund Optics, #48-605) and a reference mirror, respectively. The retroreflector and the reference mirror were mounted on a HTGM (Nutfield, QS20) and a translational stage, respectively. The light went back from the whole optical path and was separated by a second WDM, then recorded by a commercial spectrometer (Cobra-S 800, CS800-850/140, Wasatch Photonics) for NIR and a customized linear-in-k spectrometer for VIS-OCT. For noise cancellation, a fraction of the light reflected from the retroreflector and transmitted through beam splitter were coupled into a separate single mode fiber and collected by a second VIS-OCT linear-in-k spectrometer. The raw spectra in both channels are shown in supplemental Fig. 2.

We set two LED displays of 24×8 MAX7219 dot matrix modules on each side in front of eyes as the external fixation targets. The displays were controlled by the Arduino Uno R3 microcontroller with a joystick. The control program was developed in the Arduino IDE (v1.8.19). Only one dot in each display was illuminated and controlled by the moving command of the joystick. The photograph of the packaged device is shown in supplemental Fig. 3.

We used three frame grabbers (two PCIe-1437 and one PCIe-1433, National Instrument) to receive the data from three line cameras (OctoPlus, Teledyne e2v), which were triggered by the I/O device (USB-6363, National Instrument). The I/O device also controlled the electric shutter in VIS-OCT channel, 2D galvanometers and HTGM.

We developed a customized software in Microsoft Visual Studio 2019 combining with Qt (v5.15.2), QCustomPlot (v2.1.0) [46], FFTW (v3.3.5) [47], NI-DAQmx (v21.0.0), and NI-DAQmx (v21.0.0) using C/C++ for the preview and acquisition. Multi thread programing were used to acquire the data in preview and acquisition modes for the three cameras, and in parallel real-time image display in the preview. We achieved a frame rate of 20 fps (VIS) and 15 fps (NIR) with a simultaneous preview of 4 frames to show the B-scan of retina for alignment. Several imaging protocols used in this study were pre-defined and selected for acquisition.

### 4.2 Linear-in-*k* spectrometer design and configuration

Two custom built linear-in-*k* spectrometers were used for detection in the visible light channel. The spectrometer setup, similar to various near-infrared designs [22,48–50], used a diffraction grating and prism to diffract and refract the incoming light into linear-in-*k* spacing. After light collimation, the beam must pass through a diffraction grating whose incident angle (θ_i_) for the first order is calculated as:

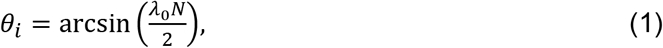

Where λ_0_ is the central wavelength and N is the number of lines per meter (1800 line/mm). The diffracted angle (θ_d_) for a given wavelength λ_m_ about the central wavelength is given by:

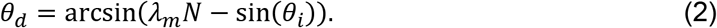

Let θ_i_ = θ_d_, and we arrived θ_i_ = θ_d_ = 28.6 degree at 532 nm per the manufacture grating specification. Since the diffracted wavelength about the center wavelength is approximately linear by wavelength, an equilateral dispersive prism is added to change the distribution. For the angle of minimum deviation, the prism incidence and deviation angles are equal. Using Snell’s Law and prism geometry, and assuming the refractive index of air is unity, the prism incident angle for the minimum deviation path (θ_pi_) at a chosen wavelength (λ_1_) is given by:

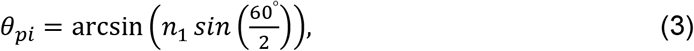

where n_1_ is the index of refraction of the prism at for λ_1_. For the center wavelength 570nm, the value for θ_pi_ = 63.4°. Using the geometry of the prism, and combining the prism dispersion with the diffraction grating, the output angle from the spectrometer can be found from the previous equations as:

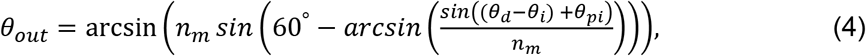

Where n_m_ is the refractive index of the prism glass at a given wavelength λ_m_. Substituting *k*_m_ = 2π/λ_m_ yields a nearly linear relationship between wavenumber and distance on the spectrometer’s camera after passing through a fixed focal length lens.

The theoretical design was first simulated in OpticStudio (Zemax) to compare the spot size of several wavelengths to the pixel size of the line camera. The spectrometer was then constructed using an 1800 line per mm volume phase holographic grating (WP-1800/532-50.8, Wasatch Photonics) and an F2 glass dispersive prism (PS852, Thorlabs). A 75 mm CA series fixed focal length lens objective (11-321, Edmund Optics) focused the dispersed light onto a 2048-pixel line camera (OctoPlus EV71Y01CCL2210-883, Teledyne e2v) with a linear-in-k distribution.

### 4.3 Per A-line noise Cancellation VIS-OCT imaging

Two spectrometers were used to provide per A-line noise cancellation. Using a beam splitter cube, part of the visible reference signal was directly transmitted to one visible light spectrometer. The other reference portion was sent to the second visible light spectrometer via a fiber coupler and wavelength division multiplexer.

After the continuous collection from two spectrometer *x_1_*(*n, τ*), *x*_2_(*p, τ*), the pixel calibration was first performed using the spectral encoding property of the excess noise of the supercontinuum [32]. The specific noise temporal patterns were used to calibrate the two spectrometers through a correlation matrix defined by the correlation of the spectrometer pixels during a shared collection period. For each pixel in *x_1_*, the most correlated pixel in x_2_ was found by locating the maximum correlation. The pixel correspondence of n and p was fitted by a second-order polynomial, which was then used to interpolate the second spectrometer data x_2_(*p, τ*) to x_2_’(*n, τ*) to align the two devices. After alignment, pixel-wise scaling factors δ(n) were calculated by δ(n) = x1(n, *τ*)/ x_2_’(*n, τ*), to match two spectra. by the calibration matrix was saved and used during processing of each image collection. The variables of the second-order polynomial and scaling factors were stored.

For noise cancellation VIS-OCT imaging, the DC components were removed by subtracting x_2_’(n) after the interpolation and scaling, followed by the regular OCT image processing methods, including digital dispersion compensation and Fourier transform. Since the spectrometer is already linear-in-k, the spectral interpolation for a linear k-spacing is not required.

### 4.4 Full-Range Wide-Field OCT with reference modulation

A high torque galvanometer scanner (Nutfield, QS20) with an attached retroreflector was used to provide full-range wide-field images. This setup was used to replace a stationary reference arm mirror and provide the modulation of the reference pathlength. Two different modulation strategies were implemented in fast and slow scanning directions.

The fast scanning modulation achieves full-range imaging. Since the intensity signal of vis-OCT is real, the Fourier transform of the signal is Hermitian, and the image has mirror terms centered on the zero-delay line. The wavenumber interferogram signals from each A-line can be represented by the signal *I*(*k, x*), where *k* is the wavenumber and *x* is the lateral A-line locations within the B-scan.

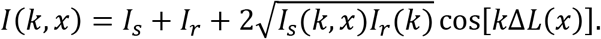

*I_s_* and *I_r_* are the intensities from the sample and reference arm respectively, and ΔL is the pathlength difference. For the sake of simplicity, we neglect *I_s_* and *I_r_* and, and only keep the third interferogram term in the following equations. Due to the nature of the retroreflector, the direction of the incoming beam is reversed without a change in angle, directly changing the path length difference. Previous full-range modulation methods typically rely on a piezo translation stage [36,51] or introducing a beam offset in the scanning mirror [52–54]. We applied a same linear ramping voltage pattern to HTGM that approximately introduce a linearly changing reference path length,

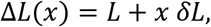

where *δL* is the pathlength increment (Unitless) introduced by HTGM. Then the modulated interferogram can be rewritten as

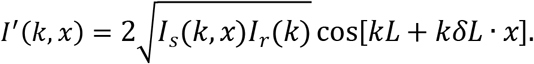

A Hilbert transform to x dimension was used to extract the complex version of the *k-x* interferogram as,

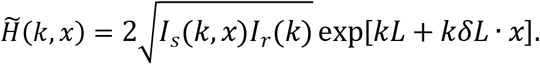

The fundamental contrast for full-range imaging is the phase modulation introduced by the reference pathlength modulation. Dense A-line sampling is required to ensure a quasi-static interferogram for an effective phase modulation. We used 1.6 um step size between A-lines, and equivalently *δL* = 0.013 (*i.e*. 13 μm reference pathlength change per 1mm fast scanning range). The pathlength modulation is implemented by feeding HTGM a same ramping voltage waveform as the fast scanning, with an amplitude scaling.

The slow scanning modulation is used to compensates the large curvature of the retina. We used an empirical non-linear function below to model the inferior part of retina along the slow scanning, assuming the fovea is at the middle B-scan frame, symmetrize it for the superior retina and applied negative voltage to the HTGM to move the image around zero-delay line despite the large curvatures.

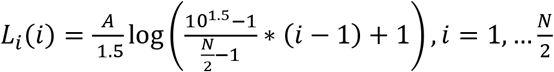

where *L_i_*(*i*) was the compensation pathlength at the *i^th^* frame. *A* was the coefficient for the compensation, equaling to −1.1. *N* was the total number of B-scan frames.

### 4.5 Imaging protocols

We prepared multiple scanning patterns and summarized them in Table 1 including preview and acquisitions of Wide Field, Full Range, High Definition, Raster, Circular-Doppler and Angio (in supplemental Fig. 4). We set the imaging line rate at 120 kHz (7.5 us exposure time). The time of all acquisition patterns was 2.18 second except Angio with 2.56 second.

**Table 1.** Imaging scan protocols.

|  | A-Scan (n) | B -scan(n) | Scanning Range (mm) | Acquisition Time (s) |
| --- | --- | --- | --- | --- |
| Preview (Radial) | 512 | 4 | 2x2 | 0.02 |
| Wide Field | 512 | 512 | 13.3*13.3 | 2.18 |
| Full Range | 8192 | 32 | 13.3*13.3 | 2.18 |
| High Definition | 2048*16 | 8 | d:6.6 | 2.18 |
| Raster | 512 | 512 | 6.6*6.6 | 2.18 |
| Circular-Doppler | 8192 | 32 | d:1.65 to d:5 | 2.18 |
| Angio | 360*2 | 384 | 3.3*3.3 | 2.56 |

The preview protocol was a 4-line radial pattern with an angular interval of π/4. Wide field covered an area of 13.3 mm × 13.3 mm with 512 A-lines × 512 B-scans. Simultaneously with HTGM, full range protocol was composed by 32 B-scans with an equal spacing, covering 13.3 mm. In each B-scan, total 8192 A-lines in 13.3 mm were scanned. High Definition protocol was to reduce speckle noise and produce high quality images [38]. In our work, we included 8-line raster and 8-line radial patterns, modulating 16 points of symmetric A-line offset along the axis which was orthogonal to the B-scan direction at each location of B-scan. The angular interval of 8-line radial pattern was π/8 and the 8-line raster cover a range of 6.6 mm with equal distance. All B-scans in High-Definition protocol have 2048 points. The range of raster protocol was 6.6 mm × 6.6 mm with 512 A-lines × 512 B-scan. The protocol of Circular-Doppler has 32 circular B-scans with equal distance, having a diameter from 1.65 mm to 5 mm. In each circular B-scan, 8192 A-lines were filled. In a square area of 3.3 mm × 3.3 mm, 360 A-lines × 384 B-scans were acquired with repeated 2 times for the Angio.

### 4.6 OCT and OCTA image processing

The raw data from both channels was processed with general procedures including DC removal, wavelength to wavenumber interpolation (only NIR), and dispersion compensation. We carried out the Fast Fourier transform (FFT) to create B-scan images by the absolute values of complex results. For per A-line noise cancellation, the DC removal used the reference spectrum acquired by the second calibrated/scaled spectrometer. For full-range VIS-OCT imaging, a Hilbert transform was performed along the lateral direction before FFT. For Doppler OCT, the angular value after FFT was extracted, and the phase difference between adjacent A-lines was calculated by subtraction. Then a 3×3 medium filter was used to remove the random noise for display.

For OCTA images, we performed the split spectrum method in NIR-OCT using 20 windows with an interval of 7.3 nm to sweep the raw spectra, then processed each swept spectra with the general procedures mentioned above and obtain the complex OCT signals. Between two adjacent complex B-scans, the axial phase group difference was corrected [55]. Then, the differences between consecutive complex B-scans at same locations created one OCTA frame.

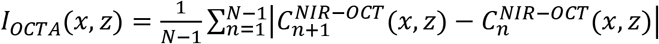

In which C(x,z) is the complex B-scan, and N = 2. In each OCTA frame, we applied the maximum intensity projection from ILM to ONL to generate the *en face* vasculature images. Finally, all these 20 *en face* images were averaged to produce the final angiography.

### 4.7 Human Imaging

All the imaging protocols have been approved by the Institutional Review Board (IRB) of Johns Hopkins University School of Medicine. The consent from volunteers was obtained before imaging.

We first moved the dual-channel light beam onto the iris of subject, then shut off the VIS-channel and used the NIR for the alignment. We asked the subjects to track the illuminated LED dot using the non-imaged eye. At the same time, four frames in the preview were continuously updated. After the desired locations of the retina were confirmed, we turned on the VIS channel, optimized the focus of the tunable lens, and started the acquisition as soon as possible.

