## Supplemental materials for "Second-generation dual-channel visible light optical coherence tomography enables wide-field, full-range, and shot-noise limited retinal imaging"

### 1. Wide field retinal VIS-OCT imaging

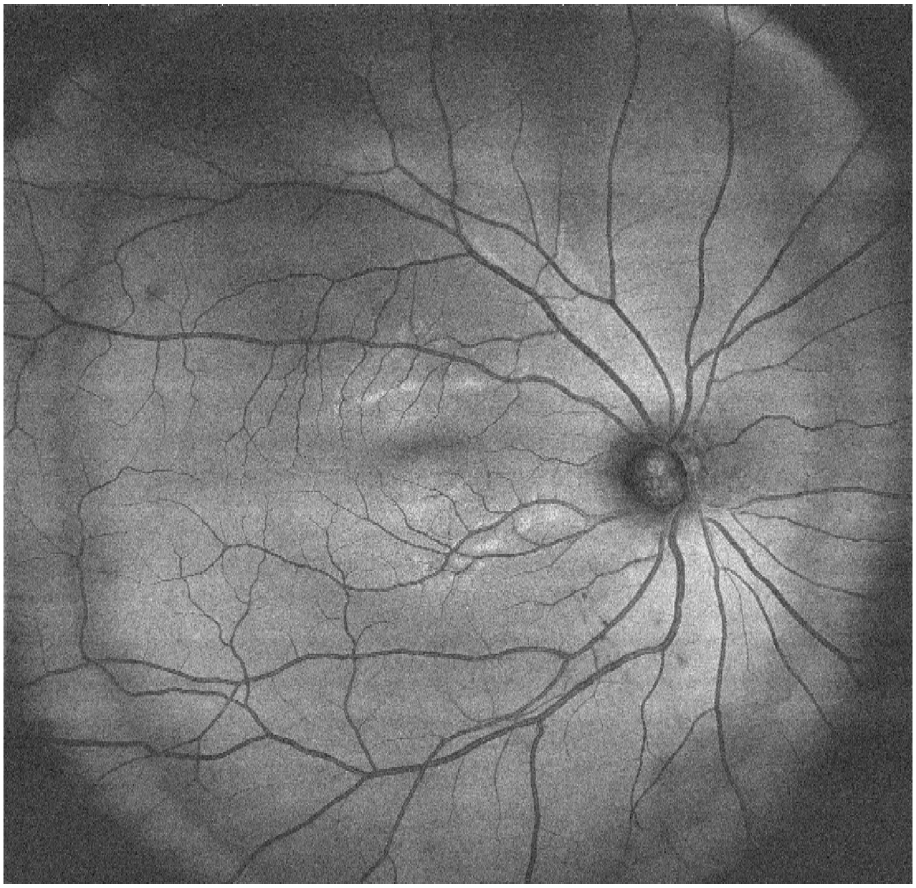

**Suppl Fig. 1**. *En face* projection of wide-field VIS-OCT imaging >60° viewing angle. The scanning density is 1024x512 acquired at 100kHz Aline rate.

### 2. Spectrum from visible and near infrared channels.

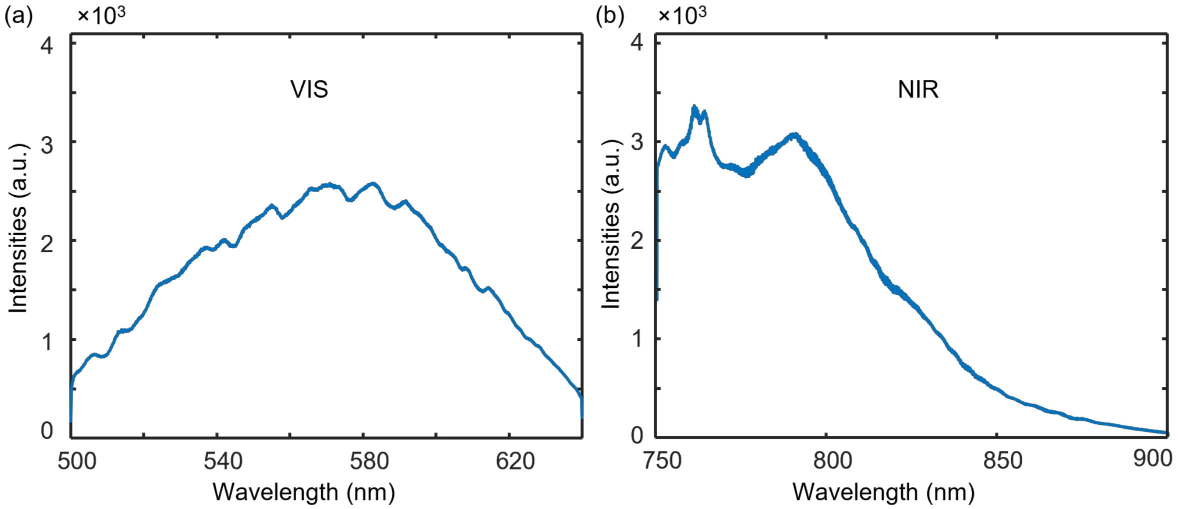


**Suppl Fig 2.** The raw spectra in VIS-OCT (a) and NIR-OCT (b) channels.

### 3. Device photograph

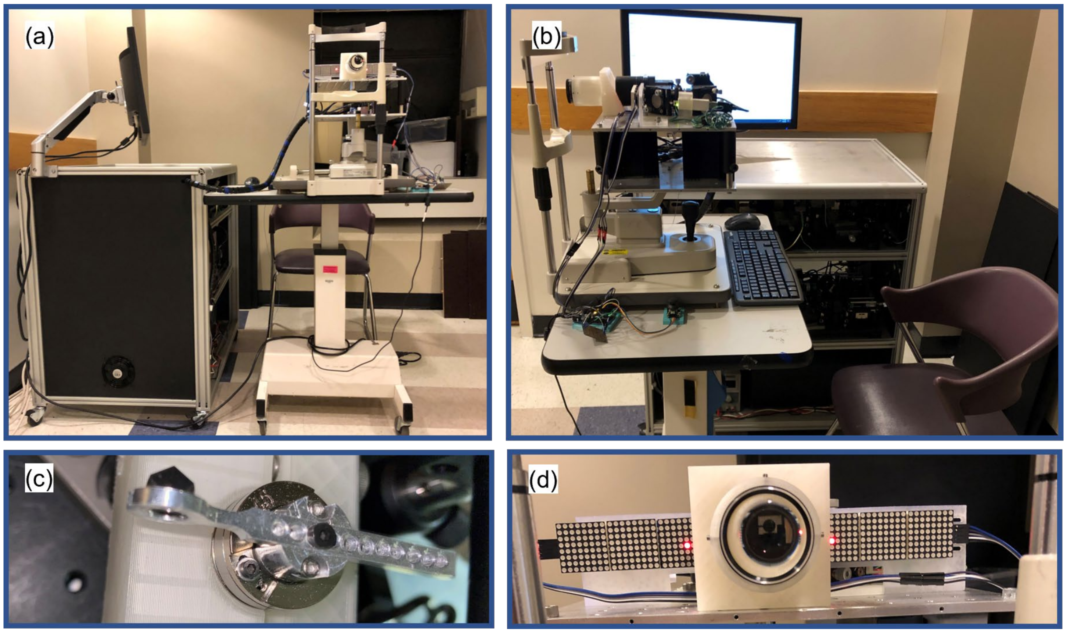


**Suppl Fig. 3.** Photograph of the device from front and side views(a-b). Photograph of custom-made mounting for the retroreflector on a high-torque galvanometer (HTGM) and the fixation target using LED arrays (c-d).

### 4. Scanning protocols

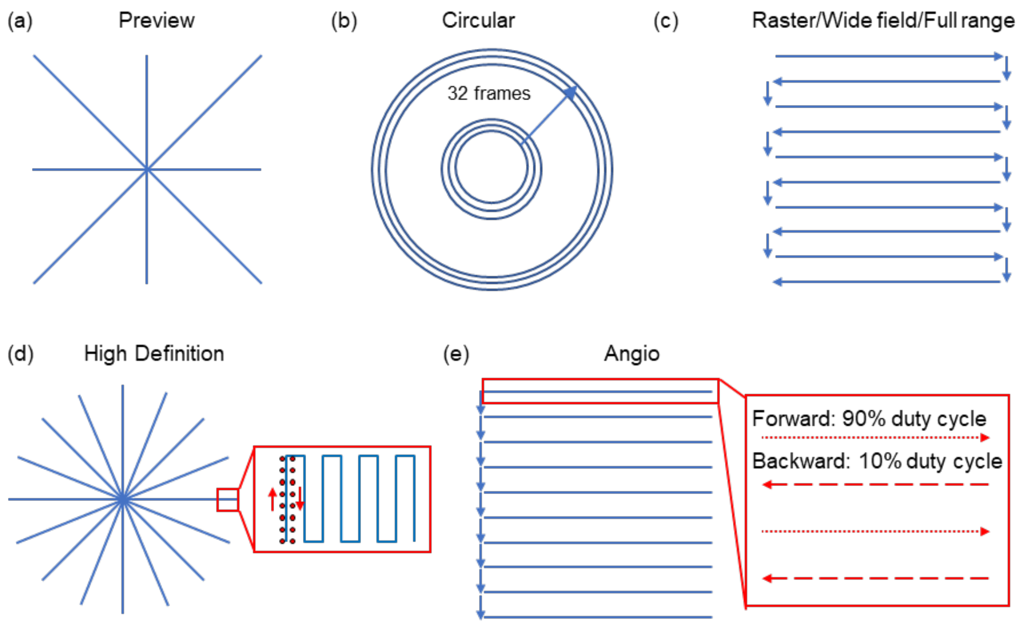


**Suppl Fig. 4**. Schematic of the scanning path for different protocols encompassed in this paper for the preview (a), circular disc scan (b), raster scan (c), high-definition scan, and (e) angiography scan.
